# Chromosome-level assembly and annotation of the grey reef shark (*Carcharhinus amblyrhynchos*) genome

**DOI:** 10.1101/2025.06.07.658398

**Authors:** Carolin Dahms, Laurent Vigliola, Leni Hiu Tong Cheung, Jerome Ho Lam Hui, Paolo Momigliano

## Abstract

To date only four of nine shark orders have nuclear reference genomes, despite next-generation sequencing advances. Particularly for threatened shark species, there is a lack of reliable genomes which are crucial in facilitating research and conservation applications. We assembled the first nuclear reference genome of the endangered grey reef shark (*Carcharhinus amblyrhynchos*) using long-read PacBio HiFi and Omni-C sequencing to reach chromosome-level contiguity (36 pseudo chromosomes; 2.9 Gbp) and high completeness (94% complete BUSCOs). BRAKER3 annotated 16,522 protein-coding genes after masking repetitive elements which accounted for 59% of the genome. We identified potential X and Y sex chromosomes on pseudo chromosomes 36 and 57, respectively. The quality and completeness of the draft genome of *C. amblyrhynchos* suggest that it has the potential to facilitate comprehensive comparative genomics, enabling researchers to investigate genetic variations and adaptations specific to this population and will help advance conservation genetic applications.

**Significance Statement:** Stemming from an ancient vertebrate lineage, sharks present an interesting evolutionary study system. A third of shark species face extinction, yet critical genomic resources necessary for research and conservation remain scarce. To address this gap, we assembled and annotated the first chromosome-level nuclear reference genome of the threatened grey reef shark (*Carcharhinus amblyrhynchos*) at high completeness. This genome will help advance studies in evolution, phylogenetics, adaptation, and conservation, offering insights not only for this species but for wider elasmobranch and vertebrate research.

## Introduction

Sharks belong to the cartilaginous fish (class Chondrichthyes), one of the oldest vertebrate lineages which emerged over 400 million years ago, encompassing a diverse array of taxa such as sharks, rays, skates and chimaeras. Having lived through five mass extinction events, sharks now face unprecedented rates of population declines due to habitat loss, overfishing and bycatch (Dulvy et al., 2014; Dulvy et al., 2021). According to the IUCN Red List, nearly one-third of shark and ray species are currently classified as threatened (Critically Endangered, Endangered, or Vulnerable; Pacoureau et al., 2021), with some families, such as Carcharhinidae (requiem sharks), exhibiting even higher proportions - up to 68% - of threatened species.

The grey reef shark (*Carcharhinus amblyrhynchos*) is one of the most abundant and widespread requiem sharks in the Indo-Pacific. *C. amblyrhynchos* has high reef fidelity (Papastamatiou et al., 2018), with rather rare long-range migrations, primarily undertaken by males (usually under 100 km; Bonnin et al., 2019). Their unique life history and habitat association has led to relatively high levels of genetic population structure compared to other shark species (Bernard et al., 2021, Cortés, 2000). Ecologically, *C. amblyrhynchos* occupies a unique niche, representing up to 50% of higher-order predator biomass on some reefs (Friedlander et al., 2014) and preying on unusually large prey for its size (Barley et al., 2020). However, life history traits such as slow growth, late sexual maturity, and long lifespan make the species particularly susceptible to overexploitation (Stevens et al., 2000). Some populations in the Indian Ocean and western Pacific have declined by more than 90% (Graham et al., 2010), raising concerns about inbreeding and loss of genetic diversity, reducing their evolutionary potential (Frankham et al., 2002) and long-term viability of remaining populations (Sherman et al., 2023).

Genomic tools are increasingly being used in conservation to monitor genetic diversity, population connectivity, and inbreeding risk (Allendorf et al., 2013). In fisheries management, these approaches have informed conservation strategies and facilitated the restoration of many commercially important fish stocks, such as Pacific salmon (Waples, 1995). The development of high-quality reference genomes is a critical step in enabling such analyses. Advances in long-read sequencing technologies, such as Pacific Biosciences (PacBio), producing lager sequence overlap and lower fragmentation (Lang et al., 2020) and chromatin conformation capture techniques like Hi-C and Omni-C, now allow for the generation of chromosome-level genome assemblies in a relatively affordable and rapid manner (Dudchenko et al., 2017).

Reliable genomes may be useful representatives of whole species to infer key conservation genomic statistics such as levels of heterozygosity, inbreeding, and demographic history. Reference genomes are key to most genetic studies commonly relying on low and mid coverage whole-genome sequences, which need to be aligned to a species-specific reference genome (Lou et al., 2021). Reference assemblies significantly improve accuracy of various other analyses such as for local adaptations and provide a crucial baseline resource for non-model species, such as genetic data deficient marine organisms (Oleksiak and Rajora, 2020).

Despite potentially severe genetic consequences of critical shark population declines, only few fisheries include assessment of population genetics (Ovenden et al., 2010), with genetic diversity and demographic history having been investigated in only about a tenth of shark species to date (Domingues et al., 2017). Contiguous high-quality genomes, allowing population and species-specific genetic assessments and genomic comparisons (Shafer et al., 2015), are currently available for less than 1% of shark species, due to lack of quality samples and their large, repetitive genomes (Pearce et al., 2021). Within the Carcharhinidae, only two nuclear reference genomes exist to date: the Oceanic whitetip shark (*Carcharhinus longimanus*; Feldheim and Pirro, 2023, unpublished) and the lemon shark (*Negaprion brevirostris*; Baeza et al., 2024).

Nevertheless, high-throughput sequencing and genome assembly projects like the Squalomix project (https://github.com/Squalomix/info/) has produced increasing numbers of shark and ray genomes (Lee et al., 2025; Baeza et al., 2024; Majeur et al., 2024; Stanhope et al., 2023; Yamaguchi et al., 2023) and drastically expanded comparative genomics (Kuraku et al., 2021). Due to their antiquity, Chondrichthyes are an interesting study system on the origins and evolution of genes (Marra et al., 2019), as well as a range of morphologies and life histories traits such as jaws (Yu et al., 2008), the cerebellum (Sugahara et al., 2017), and oviparity (Nakaya et al., 2020). Sex determination in elasmobranchs has been relatively poorly understood compared to other clades (Yamaguchi et al., 2023). Most elasmobranchs investigated so far have highly differentiated XY chromosomes thought to be of common ancient origins and lacking global dosage compensation (Wu et al., 2023), although other systems such as XX/XO in Potamotrygon sp. (De Souza Valentim et al., 2013) and possibly ZZ/ZW in *Hypanus americanus* (Schwartz & Maddock, 2002) might be confirmed as more Chondrichthyan karyotypes are investigated. To address the pressing need for shark genomic resources, we present the first assembled and annotated nuclear reference genome of the grey reef shark and identify potential sex chromosomes, which will hopefully aid conservation applications and wider evolutionary research.

## Results and Discussion

### Assembly

Three PacBio Revio Cells produced 90 Gb of sequence in 7,864,402 reads. The primary contig assembly generated by hifiasm contained 2,276 contigs with an N50 6.3 Mb, an L50 of 126 contigs with 44% GC content after purging of duplicates (Supplementary Table S1). The genome size estimate based on 21 kmers by Genomescope of 2.67 Gb from the contig assembly (Supplementary Figure S1) was similar to the final scaffolded assembly size of 2.9 Gb.

The Omni-C library used for scaffolding the contigs produced 511,721,951 read pairs of which 68.13% were high quality pairs, with 31.66% of pairs mapping >10 kb apart. 23.57% were low quality, and 8.3% unmapped. After duplicate (24.14%) removal, Omni-C sequences amounted to a coverage of 23.5X. The proximity ligation libraries greatly improved the assembly quality, as YaHS joined the contigs into 1,520 chromosome-level scaffolds (Supplementary Table S1), with an increased N50 of 90.6 Mb spanning 12 scaffolds, while 90% of the genome was covered in 36 pseudochromosomes (N95 = 276 Mb; L95 = 94; Fig. 1). Although the chromosome number of *C. amblyrhynchos* is currently not known, a haploid chromosome number of 36 is close to the expected range (n = 38-45) for *Carcharhinidae* according to karyotype studies (Stingo and Rocco, 2001) and the same as in *Triakis scyllia* belonging to the *Carcharhiniformes* (Asahida et al., 1989; Uno et al., 2020).

**Figure 1:**
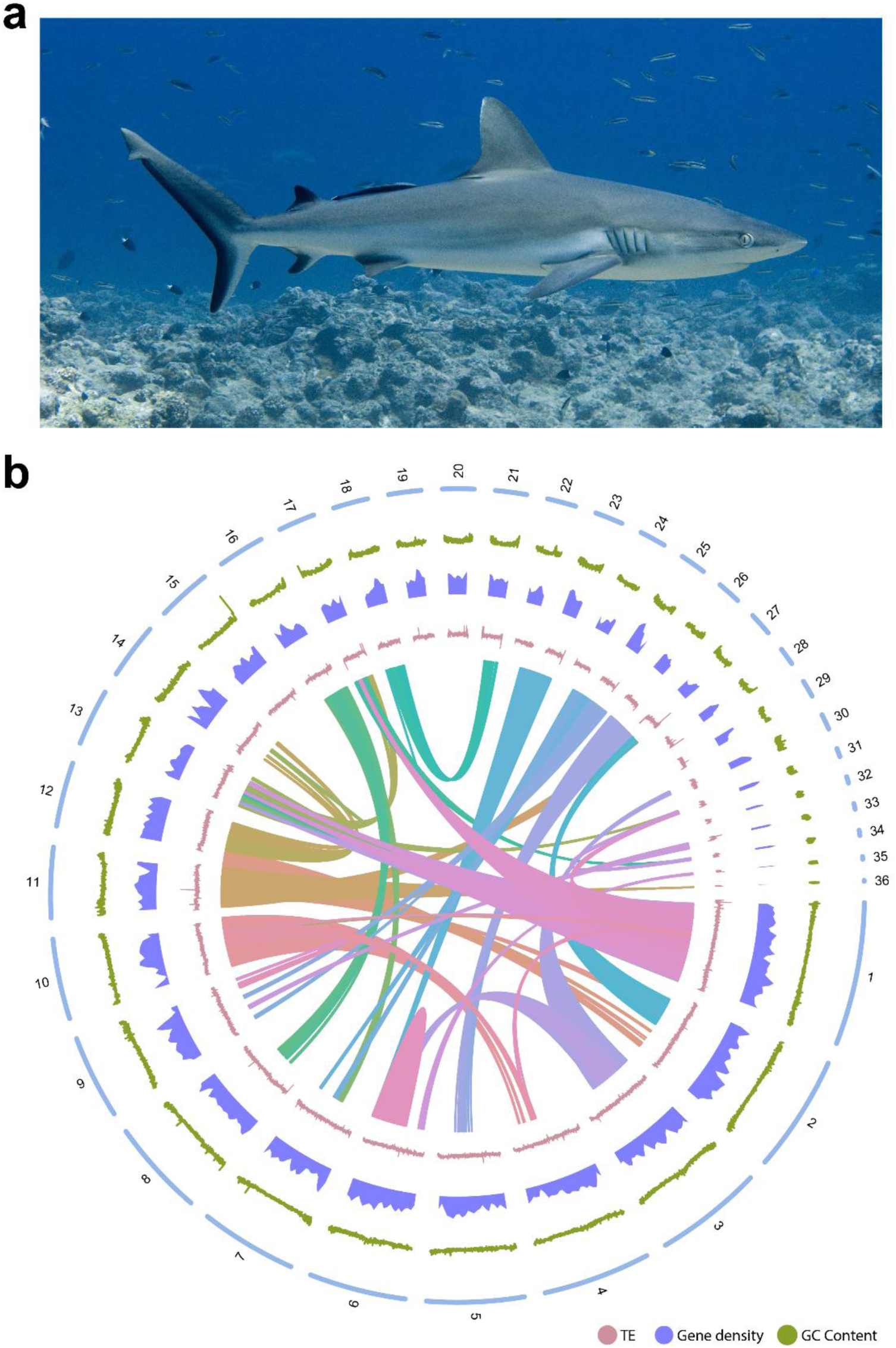
The grey reef shark and its genome assembly summary. (a) The grey reef shark (Carcharhinus amblyrhynchos), photo by Paolo Momigliano. (b) From outer to inner circles: 36 pseudo chromosomes; GC content in 100kb non-overlapping windows (green); gene density in 100kb non-overlapping windows (purple); repetitive content (pink); inner circle shows genes with conserved order between pseudo chromosomes. Alt text: The figure shows and image of a grey reef shark, and a circular plot depicting summary statistics along the main chromosomes, including gene density, repetitive content and GC content, as well as a synteny plot showing collinear gene blocks among chromosomes.

BUSCO results indicated high degree of complete BUSCOS when compared to the vertebrata lineage dataset, finding 93.6% of the 3354 genes, while 3.8% were missing, and 2.6% fragmented (Supplementary Table S2, Supplementary Figure S2). A generated Omni-C density map of the final scaffolded assembly indicated high contiguity (Supplementary Figure S3).

### Annotation

The repeat content identified with RepeatMasker amounted to 1.73Gb (59.33% of the genome) and was rich in long interspersed nuclear element (LINE) retrotransposons, comprising about 45% of the genome, the majority of which were of the L2, CR1 and Rex type (Supplementary Figure S3, Supplementary Table S3). According to sequence divergence, LINEs experienced recent expansion waves at around 7-8% and 10-13% divergence (Supplementary Figure S4b). LTRs covered 5.88%, while only 1.58% were DNA transposons, of which the Tourist and Harbinger107474 types were the most common (Supplementary Figure S4). The proportion of repetitive regions is within the expected range from 33% to 68% among sharks (Supplementary Table S2). Overall, our results are in agreement with previously published studies showing that the repeat content of sharks is rich in LINE retrotransposons, which have the highest TE activity in more recent times (Supplementary Figure S4b; Hara et al., 2018; Marra et al., 2019; Zhang et al., 2020; Tan et al., 2021; Stanhope et al., 2023).

BRAKER3 inferred 21,372 proteins encoded by 16,522 genes with an average length of 52,647 bp (ranging 7 to 2,321,391 bp; Supplementary Table S4). BLAST hits against the Uniprot SwissProt database functionally annotated 88% (14,602) of the dataset. This annotation yielded a relatively low number of predicted genes compared to related species annotations (Supplementary Table S2), due to the sub-optimal amount of RNA-seq evidence provided for model training. As obtaining tissue for endangered sharks is difficult, we had to rely on limited published RNA-seq from a different grey reef shark individual. A satisfactory, although slightly lower BUSCO completeness score of 86.5% also suggested this annotation to be rather preliminary, which we hope will provide a starting reference point for future studies.

### Sex chromosome identification

Cytogenetic studies have identified heteromorphic XY sex chromosomes in sharks, with the Y to be around one-third of the size of the X (Uno et al., 2020). In line with the expectation that the X chromosome should have reduced male sequencing coverage, while the Y chromosome should have low female coverage, we identified a 2-fold female to male depth ratio on pseudo chromosome 36 (17.3 Mb) consistent with its hemizygous state for all six males and females (Figure 2a, Supplementary Figure S5). The smaller pseudo chromosome 57 (1.5 Mb) exhibited both near 0 f/m coverage and depth ratios (Figure 2a). Thus, we can assume a XY sex determination system in the grey reef shark with 36 to be the X chromosome (Figure 2b) and 57 the Y chromosome (Figure 2c).

**Figure 2:**
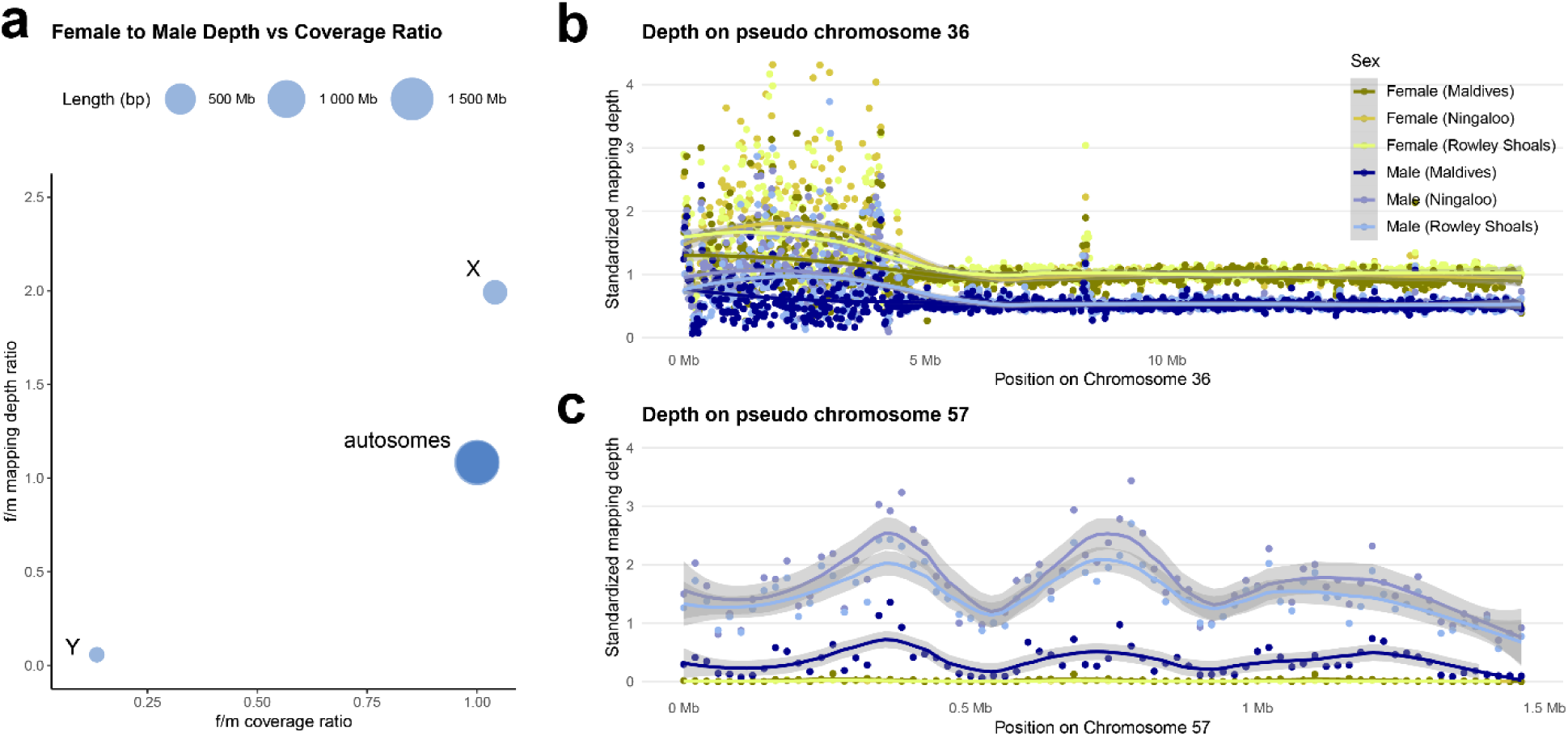
Sex identification from mapping depths and coverages. (a) Female to male sequencing coverage and mapping depth ratios for six males and females from Maldives, Ningaloo and Rowley Shoals, with near 0 coverage on the putative Y chromosome and two-fold depth ration on the X chromosome expected in a XY system; mapping depth, standardized by whole genome depth in 20kb non-overlapping windows, for the 18 individuals on pseudo chromosome 36 (b) and pseudo chromosome 57 (c). Alt text: The figure illustrates sex chromosome identification by comparing sequencing coverage and read depths from male and female grey reef sharks.

## Materials and Methods

### Library Preparation and Sequencing

PacBio and Omni-C genomic sequences were generated from dorsal fin tissue of two different male grey reef sharks, one from New Caledonia, Noumea, and one from the Southern Maldives, respectively, as tissue from the first individual did not provide enough high molecular weight DNA to cover both sequencing library preparations. The New New Caledonian individual was caught in 2016 as per Boussarie et al. (2022) under a permit from the Southern Province of New Caledonia (permits no. 479-2016/ARR/DENV and no. 2093-2016/ARR/DENV); the individual from the Maldives was obtained by this study in the Gaafu Dhaalu Atoll in 2024 under a permit from the Ministry of Fisheries, Marine Resources and Agriculture of the Maldives (permit no. NRP2023/44). Genomic DNA was extracted from alcohol-preserved tissue samples using a salting out protocol as per Momigliano et al., 2017. PacBio library preparation and sequencing was performed by the Centre for PanorOmic Sciences (Hong Kong). Briefly, quantity and integrity of the DNA sample was confirmed by Qubit and Pulse field gel electrophoresis before being fragmented to appropriate sizes. DNA fragments were damage repaired, end-repaired, and A-tailed. The SMRTbell library was produced by ligating universal hairpin adapters onto double-stranded DNA fragments. The library was checked with Qubit for quantification and Femto pulse for size distribution detection. Quantified libraries were pooled and sequenced on PacBio Revio system in 3 Revio Cells amounting to around 4 million paired-end reads containing 96 Gb of raw data.

Omni-C libraries with 570 bp inserts were prepared with the Dovetail Omni-C kit and Omni-C proximity Ligation Assay (v. 1.0, Dovetail Genomics, Scotts Valley, USA) according to the manufacturer’s protocol and sequenced on an Illumina Novaseq 6000 PE150 (Novogene, Hong Kong) as pair-end 2 × 150 bp runs, generating 153.5 Gb of raw data.

### Genome assembly

We assembled the grey reef shark genome following the Vertebrate Genomes Project (VGP) assembly pipeline (Larivière et al. 2024). Remnant adapter sequences from the PacBio HiFi dataset were removed with BBTools (Bushnell; BBDuk with the parameters *ktrim=r k=23 mink=11 hdist=1 tpe tbo*). Reads were assembled into haplotype assemblies of primary and alternate contigs using hifiasm (v.0.19.8; Cheng et al., 2021) with light purging (*-l 1 -k 21*). Duplications, repeats and contig overlaps were removed with purge dups (v.1.2.5; Guan et al., 2020) and minimap2 (v.2.26; Li 2018) using presets for HiFi data (-x map-hifi) from the primary assembly and added to the alternate assembly.

### Scaffolding

FastQC confirmed high quality of fastq reads which were mapped separately with bwa mem (Li, 2013) according to the Omni-C pipeline by Dovetail (https://omni-c.readthedocs.io/en/latest/index.html).

Valid Omni-C ligation junctions were identified using pairtools (v. 1.1.0; Open2C et al., 2024), parsed with a minimal MAPQ of “5unique” for reporting unrescuable walks, and a maximum inter alignment gap of 30. Pairs were sorted in scaffold order, and marked duplicates removed from downstream analyses. The assembly genome was scaffolded with YaHS (v.1.2.; Zhou et al., 2023) with presets. Generated Omni-C contacts with Juicer (v.1.9.9; Lander et al., 2016) were visualized in JuiceBox and checked for major misassemblies. None of the contigs or scaffolds were contaminated with either adapter or foreign species sequences according to a NCBI Foreign Contamination Screen (FCS; Astashyn et al., 2024).

### Genome Annotation

De novo repeat element libraries were built with RepeatModeler (v.2.0.5; Flynn et al., 2020) and LTRStruct (v.2.9.5) for the scaffolded assembly (from contigs > 500kb) and six other shark species (See Supplementary Table S6 for accession numbers). The program repclassifier (v.1.1; https://github.com/darencard/GenomeAnnotation/blob/master/repclassifier) was run iteratively using newly annotated known elements from our reference assembly to classify more unknown elements in three rounds; one round with repeat families from the curated Dfam database (v.3.8; Storer et al., 2020) for ancestors of *C. amblyrhynchos* (72 elements), a second round against newly classified known elements (353 additional classified elements), a third round against known elements from a consensus library of the six related shark species, which added another 213 known elements.

The genome was serially annotated and masked with RepeatMasker (v.4.1.6, Smit et al., 2015) in four rounds: simple repeats (*-a -e ncbi -noint -xsmall*); known repeats from the curated *C. amblyrhynchos* Dfam database; de novo grey repeats from reference species identified by RepeatModeler; unknown grey reef shark specific repeats from RepeatModeler (*-a -e ncbi -nolow*). The combined repeat libraries were then used to mask the genome and repeat compositions from combined analysis of all RepeatMasker rounds were summarized with ProcessRepeats. For gene prediction, the parts of genome sequences detected as repeats are soft-masked with the options *-nolow -xsmall*.

Genes were predicted in BRAKER3 (version 3.0.8; Stanke et al., 2008) with GeneMark-ETP (Gabriel et al., 2024) using AUGUSTUS v.3.3.3 trained models with RNA and protein evidence. Since we did not perform RNA sequencing, we used published *C. amblyrchynhos* retina RNA-seq from NCBI (accession number SRR2146929). The protein evidence consisted of 374 Carcharhinidae proteins (NCBI search: (txid7805[organism:exp]) NOT mitochondrion), proteins from other shark species *Rhincodon typus* (36,827), *Chiloscyllium punctatum* (33,501), *Scyliorhinus torazame* (27,605), accessed from NCBI October 2024; the well annotated *Callorhinchus milii* (GeneBank accession GCF_000165045.2) and the OrthoDB Vertebrata database (Kuznetsov et al., 2023). Genes were functionally annotated with AGAT (v.1.4.1; Dainat, 2024) based on BLAST hits against the UnipotKB/SwissProt database (accessed February 2025). Collinear gene blocks with conserved order between chromosomes were identified with MCScanX (v.1.0, Wang et al., 2012) using default parameters.

### Genome size estimation and quality assessment

k-mer counts from the PacBio HiFi reads were generated with meryl for k=18,19,20,21,22,23,31 (https://github.com/marbl/meryl). We then applied GenomeScope 2.0 (Ranallo-Benavidez et al. 2020) to the k-mer databases to estimate genome features including genome size, heterozygosity, and repeat content, with k=21 producing the best assembly (Supplementary Figure S1). Genome quality and completeness were assessed with BUSCO (v.5.6.1; Manni et al. 2021) using the vertebrate and the actinopterygii ortholog databases, with the vertebrate database providing better results. We evaluated base level accuracy (QV) and k-mers using the previously generated meryl database and merqury (version 1.0; Rhie et al. 2021) generating spectral plots which confirmed successful removal of false duplicates (Supplementary Figure S6). Assembly statistics were calculated with gfastats (v1.3.6; Formenti et al., 2022) and QUAST (v.5.2.0; Mikheenko et al., 2023).

### Identification of sex chromosomes

Sex chromosome identification was performed using whole genome sequences from six male and six female grey reef sharks from three populations, the Southern Maldives (Gaafu Dhaalu atoll, see previous methods), Ningaloo and Rowley Shoals reefs in Australia. The Australian samples were collected in 2013 and 2014 as described by Momigliano et al. (2015) under a permit from the Western Australia Department of Environment and Conservation (permit number: CE003632). DNA was extracted using a salting out protocol as per Momigliano et al., 2017 and sequenced at 10X by Novogene, Hong Kong SAR on a NovaSeq X Plus PE150 sequencing platform (see Supplementary Table S4 for sample information). Reads were mapped to the reference assembly using bwa mem (v.0.7.17), samtools (v. 1.16.1; Danecek et al., 2021) and GATK for indel realignment (v.3.8; McKenna et al., 2010). Read depths and coverages across chromosomes and average female-to-male ratios were calculated with bamdst (https://github.com/shiquan/bamdst) in 20 kb windows. A read depth ratio of 2 and coverage ratio of 1 was expected to indicate the presence of a X chromosome, consistent with the hemizygous state of the X, while female-to-male coverage and depth ratios of near 0 would be expected for the Y chromosome. For the potential sex chromosomes read depths standardized by whole genome depth were plotted.

## Data Availability

Data will be made publicly available upon publication. The final genome assembly has been deposited at DDBJ/ENA/GenBank under accession PRJEB89872. HiFi PacBio and Hi-C Illumina reads have been deposited under accessions SAMEA118362358 and SAMEA118362359, respectively. Whole genome sequences are deposited under accession PRJEB90057. The genome annotation is available at Zenodo (https://doi.org/10.5281/zenodo.15599240). Code to generate the scaffolded assembly can be found on Github (https://github.com/carolindahms/Assembly-with-Omni-C).

## Acknowledgements

The authors thank Dr Paolo Francini for his advice on genome annotation. Thanks to Sudeshna Chakraborty and Sean Law for assistance with DNA extractions and Omni-C library preparation. This work was supported by the Hong Kong Research Grant Council Collaborative Research Fund (C4015-20EF) and the Hong Kong PhD Fellowship Scheme (HKPFS).

