## Supplemental Materials for "Chromosome-level assembly and annotation of the grey reef shark (*Carcharhinus amblyrhynchos*) genome"


### Supplementary Figures

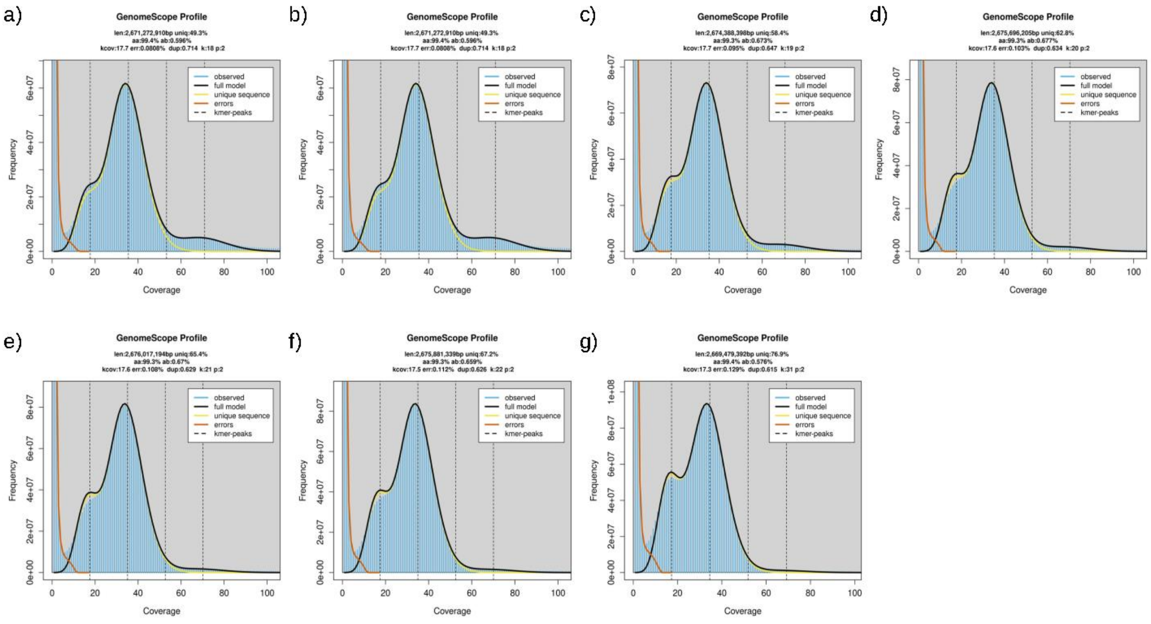


***Supplementary Figure S1: GenomeScope 2.0 profiles for kmers 17-22 and 31****. Genome size, heterozygosity and duplication rates are reported for different kmers counted with meryl. The best kmer count was 21 according to the merqury best_k script. The first peak at coverage 20X corresponds to the heterozygous peak, the second peak at coverage 40X is the homozygous peak.*

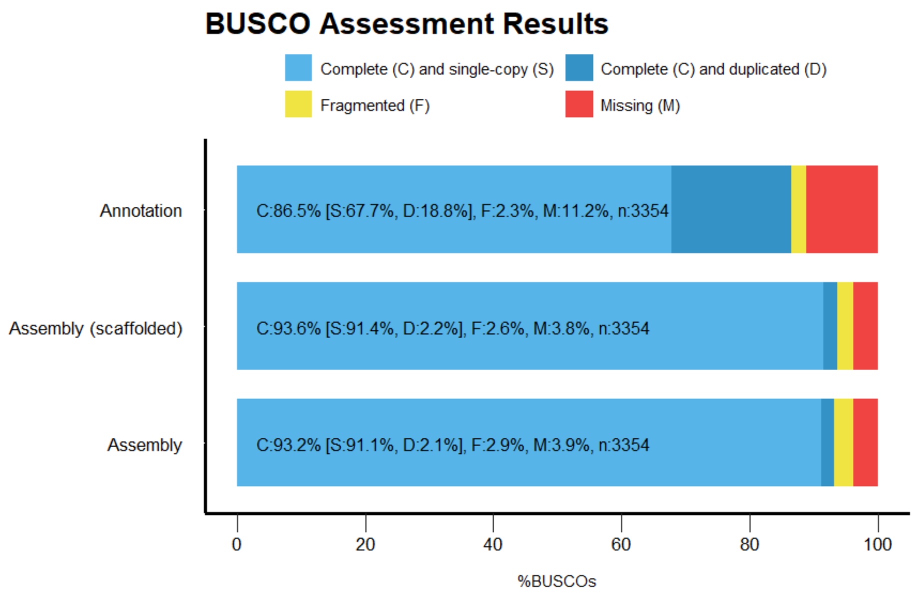


***Supplementary Figure S2: Benchmarking Universal Single-Copy Orthologs (BUSCO) results.*** *Compared are results for the primary contig assembly, the scaffolded primary assembly and its annotated version, using the vertebrata lineage data set (vertebrata_odb10, containing 3354 BUSCOs). An increase of duplicates (18.8%, Supplementary Figure S3) in the annotation is expected due to the presence of alternative isoforms in annotated gene sets (Manni et al., 2021).*

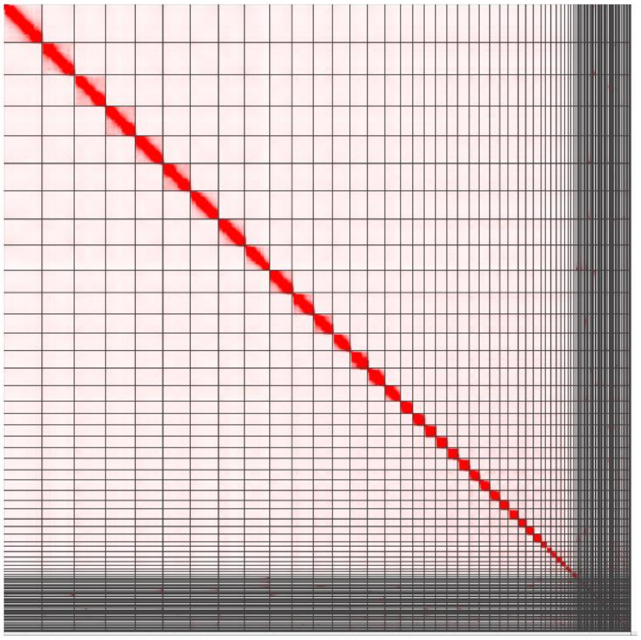

***Supplementary Figure S3: Contact map of Omni-C scaffolded final assembly,*** *showing the contiguity of the genome assembly. Scaffolds are organized from largest to smallest (left to right and upper to lower). The x and y axes indicate mapping positions of the first and second read in the read pair respectively. The colour shows the Omni-C density at each point on a log-scale. The contact map was generated with Juicebox (v.1.9.8).*

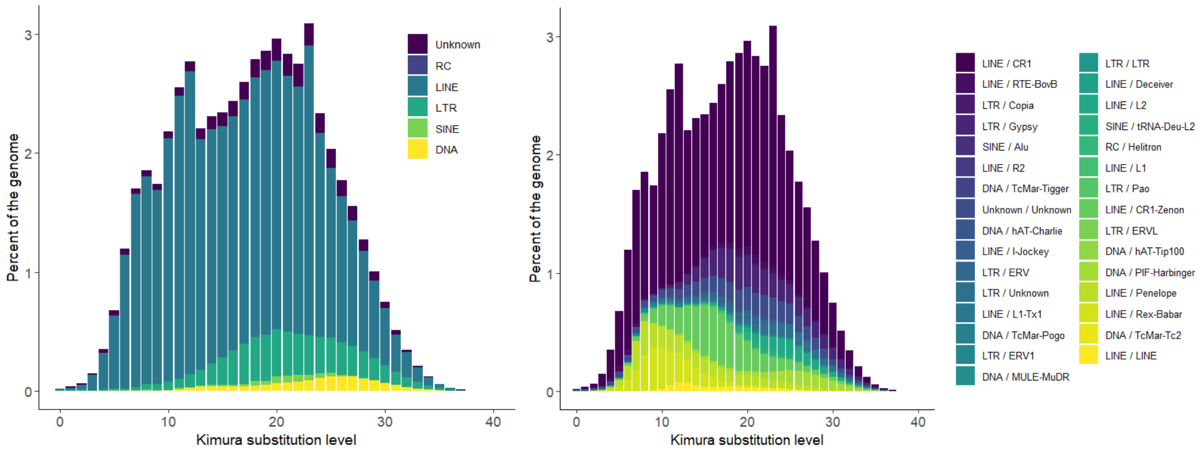


***Supplementary Figure S4: Transposable Elements (TE) landscape.*** *Copy-divergence analysis of TE classes (a) and families (b), based on Kimura distances. Proportions (%) of the genome represented by TEs are ordered according to Kimura substitution levels (arbitrary values from 0 to 40). Copies of more recent divergence are clustered on the left side of the x axis.*

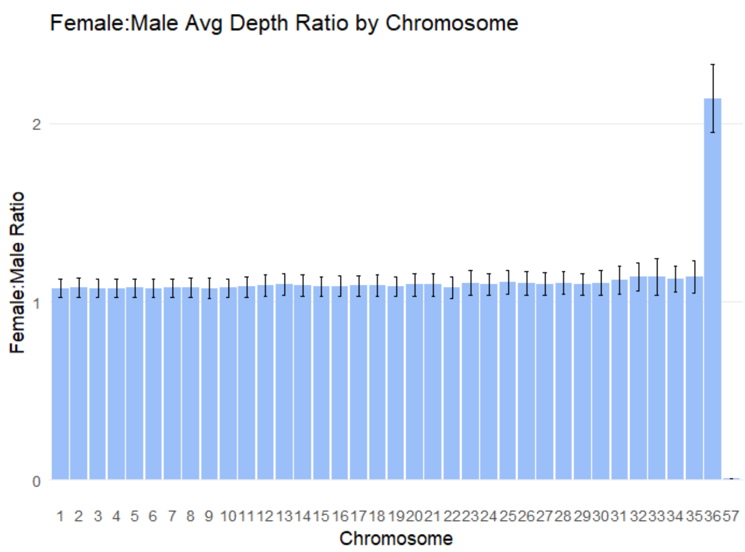


***Supplementary Figure S5: Mean Female to male sequencing depth rations across autosomes*** *(1-35) and the putative X (36) and Y (57) sex chromosomes across 18 males and 18 females from three populations from the Maldives, Ningaloo and Rowley Shoals reefs; bars represent standard errors.*

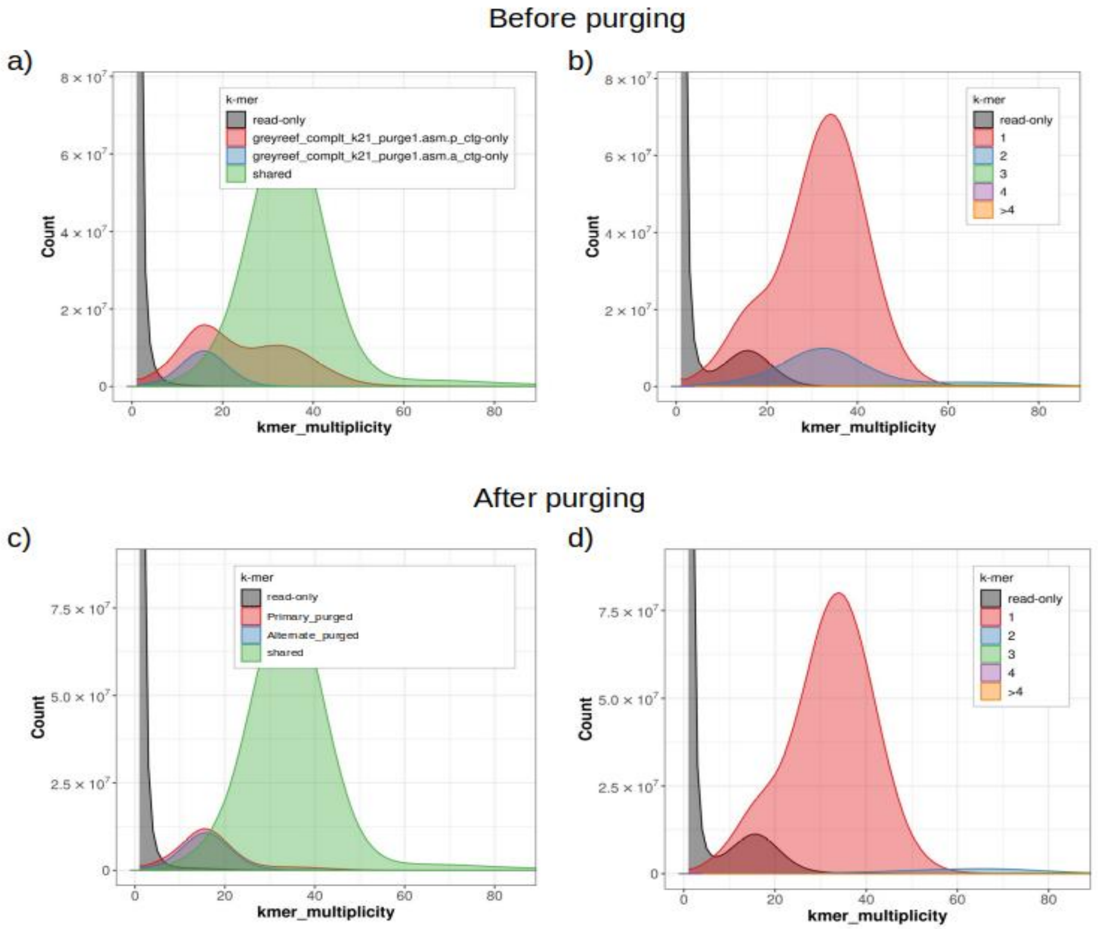


***Supplementary Figure S6: kmer plots before and after purging duplicates from the assemblies.*** *Duplicate purging greatly improved both primary and alternate assemblies, as homozygous counts represented twice in the primary assembly (indicated by high counts at diploid coverage (~38) a, red) were correctly split between assemblies (c). Reconciliation is also shown in primary-only assembly plots, where the 2-copy peak (b, blue) disappeared after duplicate removal (d). False duplications of heterozygous regions that have both their alternate alleles represented in the primary assembly, indicated by a larger primary-only peak (in red) than the alternate-only peak (in blue) at haploid coverage (~18, a) were also removed as both haploid peaks were of equal sizes (c).*

### Tables

***Supplementary Table S1: Assembly statistics*** *from primary and alternate contigs, and the scaffolded assembly from the primary contig. For the scaffolded genome, scaffold statistics are reported. Calculated with QUAST at 5kb resolution.*

| | **Primary contig** | **Alternate contig** | **Scaffolded assembly** |
|---|---|---|---|
| **Contigs/scaffolds:** | 2,276 | 2,286 | 1,520 |
| **Total contig/scaffold length:** | 2,923,473,132 | 2,927,756,164 | 2,923,561,432 |
| **Contig/scaffold N50:** | 6,200,021 | 6,139,689 | 90,619,553 |
| **Contig/scaffold L50:** | 126 | 127 | 12 |
| **Contig/scaffold N90:** | 752,330 | 736,705 | 17,339,487 |
| **Contig/scaffold L90:** | 644 | 653 | 36 |
| **Largest contig/scaffold:** | 70,791,062 | 70,791,062 | 181,693,455 |
| **Smallest contig/scaffold:** | 9996 | 8036 | 1000 |
| **GC content** | 43.98% | 44% | 43.98% |
| **Gaps** | - | - | 883 |
| **Average gap length** | - | - | 1000 |

***Supplementary Table S2: Comparison to other shark genomes published to date.*** *Shown are Sequence lengths (Gbp), number of scaffolds, N50 (of chromosomes or scaffolds), BUSCO completeness scores (%), Repetitive elements (%), predicted genes and sequencing technology used. Genome size and diploid chromosome size (2n) are estimated from molecular methods such as flow cytometry or densitometry. (1) Mirsky and Ris, 1951. (2) Hinegardner, 1976. (3) Hardie and Hebert, 2004. (4) Uno et al., 2020. (5) Schwartz and Maddock, 2002. (6) Stingo et al., 1989. (7) Schwartz and Maddock, 1986. (8) Kadota et al., 2023. (9) Stingo, 1979. (10) Stingo and Rocco, 2011. (11) Asahida, 1995. *Unpublished.*

| Family | Species | Common name | Genome size (Gbp) | Length (Gb) | 2n | Scaffolds | RE | N50 (Mb) | BUSCO | Genes | Sequencing tech | Citation |
|---|---|---|---|---|---|---|---|---|---|---|---|---|
| **Carcharhinidae** | *Carcharhinus longimanus* | Oceanic whitetip shark | 3.26[1] | 2.68 | - | - | - | - | - | - | | Feldheim and Pirro 2023* |
| | *Negaprion brevirostris* | Lemon shark | 3.62[2] | 2.29-2.58 | - | - | 50.% | - | - | - | HiSeq X Ten | Baeza et al., 2024 |
| | ***Carcharhinus amblyrhynchos*** | **Grey reef shark** | **3.34[3]** | **2.9** | **-** | **1,520** | **59%** | **90.6** | **94%** | **16,522** | **HiFi and Omni-C** | **This study** |
| Hemiscylliidae | *Chiloscyllium plagiosum* | Whitespotted bamboo shark | 3.85 | 3.78 | 102 | | 63% | 57.9 | 96% | 19,595 | BGISEQ-500 and Hi-C | Zhang et al., 2020 |
| Hemiscylliidae | *Chiloscyllium punctatum* | Brownbanded bamboo shark | 4.73 | 3.37 | 106[4] | 280,241 | 52% | 1.96 | 97% | 34,038 | HiSeq 1500 | Hara et al, 2018 |
| | *Chiloscyllium punctatum* | | | 3.38 | 104 | 275,693 | - | 69.9 | - | - | Hi-C | Hoencamp et al., 2021 |
| **Lamnidae** | *Carcharodon carcharias* | Great white shark | 6.3[5] | 3.92 | 82[5] | 9,222 | - | 3 | - | - | HiSeq | NCBI 2018* |
| | *Carcharodon carcharias* | | | 4.63 | 84 | 718 | 58% | 169.9 | 95% | 24,520 | PacBio, NovaSeq; Hi-C, Bionano | Marra et al., 2019 |
| | *Isurus oxyrinchus* | Shortfin mako shark | 4.9[3], 6.5[6] | 4.98 | 82[7] | 5,559 | 44% | 145.5 | 91% | 27,804 | PacBio Hifi and Omni-C | Stanhope et al., 2023 |
| **Rhincodontidae** | *Rhinocodon typus* | Whale shark | 3.44 | 2.93 | 102[4] | 997,976 | 38% | 0.05 | - | 19,384 | Roche 454 and Illumina | Read et al., 2017 |
| | *Rhinocodon typus* | | | 3.2 | - | | 50% | 2.56 | 84-88% | 28,483 | HiSeq 2500m | Weber et al., 2020 |
| | *Rhinocodon typus* | | 3.75[8] | 2.88 | - | 16,776 | | 70.8 | 78% | 35,334 | HiSeq X and Hi-C | Yamaguchi et al., 2023 |
| | *Rhinocodon typus* | | 3.75 | 2.66 | - | 155,537 | 38% | 0.33 | 86% | 27,005 | Roche 454 and Illumina | Hara et al., 2018 |
| **Scyliorhinidae** | *Scyliorhinus canicula* | Small-spotted catshark | 5.53[9] | 4.22 | 62 | - | - | - | - | - | - | NCBI 2021* |

| | | | | | | | | | | | | |
|---|---|---|---|---|---|---|---|---|---|---|---|---|
| | *Scyliorhinus canicula* | | | 4.22 | 62 | 645 | 68% | 199 | 92% | 24,473 | PacBio, Genomics Chromium, Bionano Hi-C | Mayeur et al., 2024 |
| | *Scyliorhinus torazame* | Cloudy catshark | 6.67, 6.45[10] | 4.47 | 64 | 458,050 | 56% | 0.28 | 84% | 28,051 | Roche 454 and Illumina | Hara et al. 2018 |
| **Sphyrnidae** | *Sphyrna mokarran* | Great hammerhead shark | | 2.77 | 78[2], 86[11] | 1,658 | 52% | 89.8 | 94% | 26,110 | PacBio Hi-Fi, Hi-C and Omni-C | Stanhope et al., 2023 |
| **Stegostomatidae** | *Stegostoma tigrinum* | Zebra shark | 3.71[8] | 2.77 | 102[4] | 1,266 | - | 76.6 | 90% | 33,222 | HiSeq X,Hi-C | Yamaguchi et al., 2023 |

***Supplementary Table S3: Repeat content for the scaffolded genome**. SINE/LINE = Short/Long Interspersed Nuclear Repeats, LTR = Long Terminal Repeats, DNA = DNA transposons. Simple refers to small RNA, satellites, simple repeats and low complexity repeats. Ran with Dfam_3.8 repeat database, with the query species Chondrichthyes.*

| Repeat Family | Primary haplotype |
|---|---|
| SINE | 14Mb (0.47%) |
| LINE | 1.3Gb (44.92%) |
| LTR | 172Mb (5.88%) |
| DNA | 46Mb (1.58%) |
| Simple | 49Mb (1.79%) |
| Unclassified | 88Mb (3.47%) |
| Total | 1.73Gb (59.33%) |

***Supplementary Table S4: Annotation statistics from BRAKER3.***

| | |
|---|---|
| Number of genes | 16,522 |
| Mean gene length (bp) | 52,647 |
| Number of CDSs | 21,372 |
| Mean CDs length (bp) | 1,775 |
| Number of exons | 221,185 |
| Mean exon length (bp) | 171 |
| Mean number of exons/gene | 10 |
| Number of introns | 199,813 |
| Mean intron length (bp) | 6,640 |
| Number of mRNA | 21,372 |
| Mean mRNA length (bp) | 63,857 |

***Supplementary Table S5: Sample information.*** *For each sample, population, sex, samping year and location coordinates are listed, as well as number of raw reads, mapped data after mapping quality (20) filtering in Mb, fraction of PCR duplicate reads (%), average read depth, fraction of coverage above 10x (%) and average insert size. Coverage and depths were calculated with bamdst in 20kb windows (https://github.com/shiquan/bamdst).*

| Sample | Population | Sex | Raw reads | Mapped data (Mb) | PCR dups | Read depth | Cov (>10x) | Insert size | Sampling year | GPS S | GPS E |
|---|---|---|---|---|---|---|---|---|---|---|---|
| MAL4 | Maldives | Male | 222,259,768 | 32,791 | 9% | 11 | 60% | 326 | 2024 | 0.21633 | 73.1595 |
| MAL5 | Maldives | Female | 220,068,372 | 32,478 | 9% | 11 | 60% | 321 | 2024 | 0.22283 | 73.1518 |
| MAL6 | Maldives | Female | 204,707,864 | 30,241 | 7% | 10 | 52% | 286 | 2024 | 0.22283 | 73.1518 |
| MAL7 | Maldives | Male | 202,157,576 | 29,881 | 10% | 10 | 52% | 329 | 2024 | 0.21506 | 73.1452 |
| MAL8 | Maldives | Female | 201,112,312 | 29,669 | 10% | 10 | 51% | 324 | 2024 | 0.20475 | 73.1419 |
| MAL9 | Maldives | Male | 202,652,256 | 29,919 | 11% | 10 | 52% | 332 | 2024 | 0.21305 | 73.14392 |
| NIN1 | Ningaloo | Male | 222,259,768 | 32,791 | 9% | 11 | 60% | 326 | 2012 | -21.97345 | 113.908 |
| NIN2 | Ningaloo | Female | 220,068,372 | 32,478 | 9% | 11 | 60% | 321 | 2012 | -21.983 | 113.91 |
| NIN4 | Ningaloo | Female | 204,707,864 | 30,241 | 7% | 10 | 52% | 286 | 2012 | -21.9832 | 113.91 |
| NIN6 | Ningaloo | Male | 202,157,576 | 29,881 | 10% | 10 | 52% | 329 | 2012 | -23.07905 | 113.732 |
| NIN7 | Ningaloo | Female | 201,112,312 | 29,669 | 10% | 10 | 51% | 324 | 2012 | -23.1615 | 113.7455 |
| NIN8 | Ningaloo | Male | 202,652,256 | 29,919 | 11% | 10 | 52% | 332 | 2012 | -23.161 | 113.745 |
| RS04 | Rowley Shoals | Female | 208,344,064 | 30,841 | 18% | 11 | 53% | 309 | 2011 | -17.253 | 119.360 |
| RS10 | Rowley Shoals | Male | 211,716,628 | 31,181 | 6% | 11 | 54% | 244 | 2011 | -17.283 | 119.370 |
| RS74 | Rowley Shoals | Female | 244,751,340 | 36,238 | 20% | 12 | 66% | 297 | 2011 | -17.252 | 119.360 |
| RS78 | Rowley Shoals | Female | 264,643,426 | 38,971 | 43% | 13 | 66% | 280 | 2011 | -17.2837 | 119.3705 |
| RS82 | Rowley Shoals | Male | 212,160,358 | 31,388 | 19% | 11 | 54% | 299 | 2011 | -17.252 | 119.360 |
| RS96 | Rowley Shoals | Male | 186,146,446 | 27,036 | 37% | 9 | 41% | 334 | 2011 | -17.253 | 119.360 |

***Supplementary Table S6: Reference genomes and accession numbers of closely related shark species used for repeat annotation.***

| Species | GeneBank accession | Genome size | Repeats | Reference |
|---|---|---|---|---|
| Oceanic whitetip shark (*Carcharhinus longimanus*) | GCA_030264375.1 | 2.68 Gbp | - | Feldheim and Pirro, 2023 |
| Great white shark (*Carcharodon carcharias*) | GCA_003604245.1 | 4.29 Gbp | 58.5% | Marra et al., 2019 |
| Brownbanded bambooshark (*Chiloscyllium punctatum*) | GCA_003427335.1 | 3.37 Gbp | 51.87% | Hara et al., 2018 |
| Whale shark (*Rhincodon typus*) | GCA_021869965.1 | 2.93 Gbp | 38.3% | Read et al., 2017 |
| Great hammerhead (*Sphyrna mokarran*) | GCA_024679065.1 | 2.77 Gbp | - | Stanhope et al., 2023 |
| Lemon shark (*Negaprion brevirostris*) | GCA_030324005.1 | 2.29-2.58 Gbp | 64-71 % | Baeza et al., 2024 |
